# DNA metabarcoding Uncover the Diet of Subterranean Rodents in China

**DOI:** 10.1101/2021.09.20.461124

**Authors:** Xuxin Zhang, Yao Zou, Xiaoning Nan, Chongxuan Han

**Affiliations:** Key Laboratory of National Forestry and Grassland Administration on Management of Western Forest Bio-Disaster,College of Forestry,Northwest Agriculture and Forestry University,Yangling, Shaanxi, China

## Abstract

**Objective:** In the past, the zokor, which lived in northern China, caused great harm to agriculture and forestry production due to its large and sophisticated diet. Since the rat lives underground for most of its life, researchers know little about its dietary habits. Further understanding of its diet in the field is of important meaning for developing green and sustainable control strategies for the rat.

**Methods:** Longde County in Liupan Mountain area of Ningxia Hui Autonomous Region was selected as the interest area to capture zokor and investigate the species of habitat plants.We selected chloroplast *trn*L gene and eukaryotic internal transcription spacer 1 (ITS 1) primers to amplify DNA from the gastric contents of zokor,and then sequenced on Illumina Miseq PE300 platform.

**Results:** The gastric contents of *Eospalax smithii* (n=16) and E.cansus(n=9) were analyzed by operational taxonomic units (OTU) clustering and amplicon sequence variants(ASVs).The OTU clustering method obtained 2,995 OTUs, and the ASV method obtained 4,657 ASVs. The ASV method was better than the OTU clustering method, and the ASV method was adopted in the subsequent analysis. The food list of 32 families, 80 genera and 154 species was obtained by ASV method after the error was removed. The food composition of zokor was evaluated by relative abundance(%RA) method and frequency of occurrence(%ROO) method. At the Family level, it was found that zokor mainly fed on Asteraceae, Poaceae, Rosaceae, Pinaceae, Brassicaceae, Apiaceae, etc. At the Genus level, they are mainly*Echinops, Littledalea,Artemisia,Picea, Cirsium, Elymus* and so on. The diet alpha diversity of *E.cansus* was slightly higher than that of *E.smithii* (*P* > 0.05). The correlation coefficient between Sobs index of alpha diversity and body weight of zokor was −0.382 (*P* = 0.059). The diet beta diversity proved that most zokors (22/25) clustered together, with low heterogeneity. They fed positively on *Calamagrostis, Cirsium, Echinops, Medicago, Sanguisorba* and *Taraxacum*. We found that zokor mainly fed on the roots of perennial herbs(PH), which were rich in water, carbohydrate, fat and protein, which provided an important energy source for its survival.

**Conclusion:** High-throughput sequencing(HTS) based DNA metabarcoding technology has well revealed the diet of zokor, which are generalist.

## Introduction

The subterranean rodent is a small herbivorous mammal that spends its entire life under the ground [1, 2]. They generally have well-developed forelimbs suitable for digging, vision degeneration, smell and hearing developed. Underground rats need a lot of energy to dig and maintain a complex system of tunnels, especially in hard, dry soil [3]. This high energy consumption forces the underground rat to expand its food forage and become a food generalist [2, 4]. In addition, Heth analyzed the feeding behavior of moles (Spalacidae) and concluded that animals that forage underground, with large search costs and diverse food resources, must collect all edible food species found when digging burrows.This, combined with the non-directed search pattern, will produce a generalist foraging behavior [5].

Myospalacinae is a subterranean rodent endemic to East Asia [6], and mainly distributed in a wide area north of the Yangtze River Basin in China. According to our research and literature records, there are currently 9 species of zokor in China, including 6 species of *Eospalax* and 3 species of *Myospalax* [6]. The average daily food intake of *E.cansus* is 105.7g for herbaceous plants and 60.5g for tree roots, and there is a highly prominent positive correlation between daily food intake and body weight [7]. The distribution of the zokor in China also conforms to Bergmann’s rule, that is the body weight of the zokor increases with the latitude. The body weight of *E.rothschildi* living in Qinba Mountains(QBM) was markedly lower than that of *E.fontanierii, E.cansus*, and *E.smithii* living in Loess Plateau(LP). It is also significantly smaller than *M.aspalax* in the northern Mongolian Plateau(MP). At the same time, the body weight of *M.aspalax* was remarkably higher than that of the three zokors in the LP.(W_QB_ < W_LP_ < W_MP_,*P* < 0.01; **See Fig S1**). In QBM, zokor lives mainly in farmland and alpine meadow. In LP, the species and population of zokor were the highest owing to less woodland and more cultivated land. In MP, the woodland is scarce, the range of movement of zokor is large, and the harm to the grassland by feeding on grass roots is larger. Studies have shown that the zokor has caused serious losses to the industry of new woodland [8–10], crops and herbs [11, 12]. The damage of zokor is particularly serious on LP. Chinese scientists’ research on the zokor began with the prevention and control of the disaster of zokor.

The gnaw of zokor causes serious losses in new woodland and young forest. In the light of our investigation and literature records, in terracing fields without prevention and control measures, the damage rate of seedlings in new woodland can be as high as 82.5% [8]. With the deepening of the concept of ecological development, the control of zokor has shifted from pesticide poisoning and physical killing to the concept of protecting the purpose tree species [13]. The purpose of this study was to explore the diet of zokor, to understand the interaction mechanism between zokor and plant, to explore the status and function of zokor in the ecosystem, and to objectively analyze the degree of harm of zokor, so as to provide scientific basis for making effective rodent control measures [14].

At present, traditional methods are used to study the diet of zokors. For instance, captive feeding [16], observation of food accumulation in caves [21], observation of foraging behavior and microhistology of gastric contents [22,23], etc. The above methods are limited in terms of season and food diversity, and can only reflect the dietary trait of zokor in part of the time period, it is even more difficult to reveal the diet in wild zokor. Microhistology is the highest resolution technique in the current study of diet of zokor. This method requires accurate analysis of partially digested plant fragments but may overemphasize the indigestible groups [24]. However, it is still limited by the researchers’ ability of plant classification and heavy workload, and these factors make the study on eating habit of zokor not comprehensive and objective.

In recent years, with the improvement of molecular biology database and the emergence of high-throughput sequencing (HTS). DNA metabarcoding, which integrates the idea of DNA barcode and the HTS technology, is born at the historic moment. DNA metabarcoding is not restricted by prey species, and can identify prey at the taxonomic Species level, especially suitable for diet research of difficult-to observe or special habits, and allows parallel processing and sequencing of large samples [25].

In the selection of diet samples, researchers generally choose different sampling methods in line with the study species. The most common type is feces, because it is the easiest way to obtain and has the minimal damage to predators, especially some rare and mysterious species [20,24,26,27].However, due to the digestion and decomposition of animals, the food DNA in feces degrades too much. Therefore, regardless of the species, the best sample is the stomach contents, where the degradation degree of food in the stomach is the lowest and relatively more food DNA sequences can be obtained [28]. DNA metabarcoding technology mainly uses universal primers to amplify food DNA fragments of animals, so the selection of primers is particularly crucial. For herbivores, chloroplast gene fragments such as *trn*L(UAA) intron and its above P6 Loop [4, 18, 24, 29, 30], *rbc*L [31–33] and *trn*H-*psb*A [18, 46] are usually selected. Amplified the fecal DNA of herbivorous geese by *trn*H-*psb*A, *rbc*L, *mat*K and *trn*L respectively, resaechers found that the success rate of *trn*L amplification was the highest [18]. Another study suggests that the combination of three primers can improve the food variety by 30% [26]. The use of *trn*L (UAA) P6 loop combined with ITS1-F/ITS1*Poa*-R and ITS1-F/ITS1*Ast*-R can improve the species resolution of Poaceae and Asteraceae [24]. Therefore, it is extremely vital to combine different types of primers for the study of the diet of zokor, which is a broad-feeding animal.

For a long time, the food sequence analysis of metabarcoding was mainly based on the OTU clustering method. The idea was to generate operational taxon Unit (OTU) by clustering the sequencing results after amplification and sequencing of molecular markers (barcodes). Each OTU includes sequences within the sample that differ very little (usually within 3%). There is no specific classification information of OTU and it cannot be matched one-to-one with species. Simulation studies have found that the number of OTU obtained by OTU clustering analysis is often far more than the actual number of species, which indicates that OTU clustering analysis has a high number of false positives and is too dependent on reference database to be compared between different samples [34].Recently, researchers have developed new methods to resolve amplicons sequence variants(ASVs) from Illumina-scale amplicons data without imposing arbitrary distinct thresholds that define molecular OTUs [34]. Many microbiology-related studies have compared OTU clustering method with ASV method. The ASV method maintains the extensive ecological pattern observed by OTUs in the analysis of fungal diversity, including diversity sequencing, and at the same time improves the reproduceability and data sharing of fungal studies, which can reproduce the biogeographic information hidden by OTU clustering method [35].Under the condition that the sequencing depth is sufficient to capture the complexity of the community, the ASV method is superior to the OTU clustering method in terms of community diversity, especially in terms of fungal-related sequences [36]. ASV method has the strongest correlation to infer pollutants and sample biomass, and the ASV method has good sensitivity and precision, which excel traditional methods [37]. The data set of QIIME2 analyzed by ASV could identify the sample collection point more accurately after removing the genera with lower relative abundance (< 1%) [38]. In terms of intestinal microbiota of aquaculture shrimp: Compared with traditional OTU clustering methods, ASV method has practicality in finding a wider range of *vibrio* species diversity [39]. A number of recent environmental DNA studies have also chosen to use ASV methods in data processing. Such as visualization of seasonal variations in airborne pollen [40], food choices of endangered small mammals [41], and food composition of native and alien invasive fish [42]. It is a necessary attempt to use ASV method to process and analyze the data of diet sequencing.

In the Liupan Mountain of Ningxia Hui Autonomous Region, the zokor is the most harmful rodent in the area of returning farmland to forest, and the damage to new woodland and young forest is extraordinarily serious. The research on local zokor is mainly focused on prevention and control, and the basic research on the diet of zokor is still in the traditional direct observation method [8–10]. It is helpful to provide theoretical basis for the control of zokor in this region and even in China to study the diet of zokor by using the cutting-edge DNA metabarcoding technology.

## Materials and methods

### Study area

The study area is located in Longde County, Guyuan City, Ningxia Hui Autonomous Region, China (E: 106°9 ‘54.49 “; N:35°30’ 10.76 “), with an altitude of 2250m. The study area covers an area of about 300 hectares. The study area is the project area of returning farmland to forest and grassland, the main tree species are *Pinus tabuliformis, Pinus sylvestris, Picea crassifolia*, and *Hippophae rhamnoides*. The main herbaceous plants genera are *Elymus, Leymus, Artemisia, Cirsium, Bupleurum*,and so on.

### Study area flora

We investigated the flora in the study area on September 3, 2020, October 26, and April 18, 2021. The plant species were identified according to the morphological characteristics of *Flora of China* and *Flora of Ningxia*. References to the PPG I system for ferns [42], Christenhusz [43] for gymnosperms, and APG IV for angiosperms [44]. Due to the limited survey time, the common plants in the same area were also referred to in *Flora of Liupan Mountain* and *A Guide to Common Plants on the Loess Plateau*. Seven plots with an area of 1ha were set within the main activity area of zokor. The species, quantity and height of arborous layer, shrub layer and herbal layer were investigated in the plots with an area of 1ha.

### Acquisition of stomach contents of zokor

As zokor is a local pest, our samples came from a “rat hunt” by the local forestry department. We obtained the wild zokor with the approval and assist of Forest Pest Control station of Natural Resources Bureau of Longde County, Ningxia Hui Autonomous Region, and the research area is not a nature reserve. The rat is trapped in a ground arrow trap, which causes the rat to become unconscious within a short time of being hit, greatly reducing its pain. The harvesting of zokor and the collection of stomach contents was approved by the Forest Pest Control Station in Longde County. All experiments were approved and supervised by the Science Ethics Committee of Northwest A & F University. Dissection was performed after recording relevant body indexes of zokor. The digestive tract was stripped out, and the stomach contents of zokor were clamped with sterilized scissors and forceps into a 2mL cryopreservation tube. After collection, the contents were immediately stored in liquid nitrogen gas phase for subsequent experiments. Among the captured zokors, 25 were selected for diet analysis. Among them, *E.smithii*(*n*= 16; 8 male,8 female), *E.cansus*(*n* = 9; 6 male,3 female).(**Table 1**.) The average body weight of male was 315.0g (±40.7 SD, n =14) and the female zokor was 204.2g (± 21.2SD, n = 11). The body weight of the two species of zokor in the study area collected by us was analyzed (n=44). There was no significant difference in body weight between different species of zokor, and the body weight of male zokor was significantly higher than that of female (*P*<0.01) (data not shown).

**Table 1.** specimens information.

| plot ID | altitude/m | longitude&latitude | specimens ID | date | gender | weight/g |
| --- | --- | --- | --- | --- | --- | --- |
| LD-A | 2260 | 106.16 E;35.51 N | EC-1 | 2020.10.3 | female | 214.0 |
|  |  |  | EC-7 | 2020.10.25 | female | 174.5 |
|  |  |  | ES-4 | 2020.10.25 | female | 187.9 |
|  |  |  | ES-13 | 2020.10.26 | female | 197.6 |
|  |  |  | ES-14 | 2020.10.26 | female | 185.4 |
| LD-B | 2271 | 106.16 E;35.50 N | ES-5 | 2020.10.25 | male | 242.7 |
|  |  |  | ES-6 | 2020.10.25 | female | 208.2 |
|  |  |  | ES-7 | 2020.10.25 | male | 357.2 |
| LD-C | 2268 | 106.16 E;35.50 N | ES-1 | 2020.9.3 | male | 355.7 |
|  |  |  | ES-8 | 2020.10.25 | male | 307.5 |
|  |  |  | ES-9 | 2020.10.25 | female | 171.7 |
| LD-D | 2332 | 106.16 E;35.49 N | ES-2 | 2020.9.3 | male | 226.4 |
|  |  |  | ES-10 | 2020.10.25 | female | 221.4 |
|  |  |  | ES-11 | 2020.10.25 | female | 308.8 |
| LD-E | 2242 | 106.16 E;35.49 N | EC-5 | 2020.10.3 | male | 357.0 |
|  |  |  | ES-3 | 2020.9.3 | male | 267.7 |
|  |  |  | ES-12 | 2020.10.25 | female | 220.0 |
| LD-F | 2242 | 106.16 E;35.50 N | EC-2 | 2020.10.3 | male | 270.0 |
|  |  |  | EC-3 | 2020.10.3 | male | 358.0 |
|  |  |  | EC-4 | 2020.10.3 | male | 352.0 |
|  |  |  | EC-6 | 2020.10.3 | male | 327.0 |
| LD-G | 2262 | 106.19 E;35.57 N | EC-8 | 2020.10.26 | female | 226.7 |
|  |  |  | EC-9 | 2020.10.26 | male | 337.7 |
|  |  |  | ES-15 | 2020.10.26 | male | 338.1 |
|  |  |  | ES-16 | 2020.10.26 | female | 238.6 |

Note: ES: *E.smithi*; EC:*E.cansus*.

### DNA barcode selection

We amplified DNA extracted from stomach contents of zokor by *trn*L and ITS 1 primers, respectively. The sequence of *trn*L primers: *trn*L-*c*-F_*trn*L-*h*-R (*c*-F: CGAAATCGGTA-GACGCTACG; *h*-R: CCATTGAGTCTCTGCACCTATC) [18]; ITS 1: ITS 5-F_ITS 2-R: (F: GGAAGTAAAAGTCGTAACAAGG; R: GCTGCGTTCTTCATCGATGC) [26].

### DNA extraction and PCR amplification

Total DNA was extracted from stomach contents of 25 zokors according to the manufacturer’s protocol of the E.Z.N.A.^®^ soil DNA kit (Omega Bio-Tek, Norcross, GA, U.S.). 1% Agarose gel electrophoresis was used to detect the quality of DNA extraction. DNA samples were quality checked and the concentration was quantified by NanoDrop 2000 spectrophotometers (Thermo Fisher Scientific, Wilmington, DE, U.S.).

Chloroplast *trn*L-*c*-F_ *trn*L-*h*-R and eukaryotes ITS 1 primers were selected for PCR amplification. Transgen AP221-02:Transstart FastPFU DNA Polymerase,20μL reaction system: 5 × *TransStart* FastPfu buffer(4μL),2.5 mM dNTPs(2μL), forward primer(5μM) 0.8μL, reverse primer(5μM) 0.8μL, *TransStart* FastPfu polymerase 0.4μL,BSA 0.2μL, template DNA 10 ng, up ddH_2_O to 20μL, 3 replicates for each sample. PCR reaction parameters(1) *trn*L-*c*-F;*trn*L-*h*-R: 3 mins at 95°C; 35× (30 s at 95°C; 30 s at annealing temperature 56°C; 45 s at 72 °C); 10 mins at 72 °C, 10 °C until halted.(2) ITS 5-F_ ITS 2-R: 3 mins at 95°C; 35 × (25 s at 94°C; 30 s at annealing temperature 55 °C; 25 s at 56 °C); 10 mins at 72 °C, 10 °C until halted.

### Illumina sequencing of *trn*L and ITS 1 amplicon

Agarose gel electrophoresis was performed to verify the size of amplicons. Amplicons were subjected to paired-end sequencing on the Illumina MiSeq PE300 sequencing platform at Majorbio Bio-Pharm Technology Co. Ltd. (Shanghai, China). The raw reads were deposited into the NCBI Sequence Read Archive (SRA) database (Accession Number: PRJNA753876).

### Sequencing data processing

Fastp (v0.19.6) [45]was used for quality control of the original sequence, and FLASH (v1.2.7) [46] was used for splicing. Filter the reads with a tail mass value of 20 or less bases and set a window of 50bp. If the average mass value in the threshold is lower than 20, remove the rear bases from the threshold. Filter the reads with a tail mass value of 50bp and remove the reads containing N bases. According to the overlap relationship between PE reads, pairs of reads were merged into a sequence, and the minimum overlap length was 10bp. The maximum error ratio allowed in overlap area of spliced sequences was 0.2, which screened the non-conforming sequences. The samples were differentiated according to the barcode and primers at both ends of the sequence, and the sequence direction was adjusted. The number of mismatches allowed by barcode was 0, and the maximum number of primer mismatches was 2.

We used OTU clustering and ASV methods to compare the processed data and compare the differences in species annotation between the two methods. OTU clustering: bioinformation statistical analysis was performed on OTU at 97% similar level, and UPARSE7.0.1090 was used for OTU clustering. NT_V20200604 database was selected, and RDP Classifier2.11 was used for sequence classification annotation. Usearch7 was used for OTU statistics, Mothur1.30.2 was used for Alpha diversity analysis, Qiime1.9.1 was used to generate abundance tables for each taxonomic level, and the distance of beta diversity was calculated. ASV methods: Based on the default parameters, the DADA2 [48]plug-in in QIIME2 process [47] was used to de-noise the optimized sequence after quality control splicing. The representative sequences of ASV were classified and identified using the “NR/NT Collection” (https://blast) of GenBank database NCBI. https://blast.ncbi.nlm.nih.gov/Blast.cgi. Taxonomic annotation of ASV in the “Nucleotide Collection (NR/NT)” of GenBank database NCBI was performed using the multi_blast method in QIIME2 (V2020.2). Sequence consistency: 0.8 Sequence coverage: 0.8. Through the diversity cloud analysis platform (QIIME2 process) of Majorbio Bio-Pharm Technology Co. Ltd. for subsequent data analysis. Two indexes, %RA (Relative abundance) and %FOO (Frequency of Occurrence) [20, 27], were used to evaluate the food composition of zokors. % RA refers to the percentage of the occurrence frequency of the sequence number (ASVs) of a certain food species in the total occurrence frequency of all food ASVs. The calculation formula is as follows: % RO_i_ = (N_i_/∑N_i_) ×100%; ∑N_i_ is the sum of the ASV occurrence times of all the food of the animal. %FOO refers to the percentage of the samples containing ASV of a certain food in the total number of samples, and the calculation formula is: %FOO_i_= (N_i_/N) ×100%, where N_i_ represents the number of samples with ASV of class i food, and N is the number of effective samples [27].

## Result

### Plant species in the study area

We surveyed the flora in the study area in September 2020, October 2020 and April 2021, respectively. In the study area, after the policy of returning farmland to forest and grass was implemented, the main tree species planted were *Picea crassifolia, Pinus sylvestris, Larix gmelini, Betula platyphylla, Armeniaca sibirica* and *Populus davidiana*. Common shrubs are *Hippophae rhamnoides* and *Sambucus adnata*, etc. The common herbs are mainly weeds such as Asteraceae, Poaceae, Fabaceae, Rosaceae and Apiaceae. See **Appendix 1** for detailed plant species.

### Sequencing information statistics

HTS showed that effective food DNA was amplified from all gastric content samples. We used the traditional OTU clustering method to obtain 2,694,529 sequences. After denoising, 1,986,045 sequences were obtained, and 2,995 OTUs were obtained by clustering. 99.80% OTU annotated as Eukaryota. In Eukaryota’s sequence, Fungi accounted for 67.01%, Viridiplantae 31.32%, Metazoa 1.62%, and unclassified 0.05%. We also used the ASV method to obtain 3,514,769 sequences, and 1,852,644 valid sequences were obtained by DADA2 denoising and 4,657 ASVs. 99.81% of the effective ASV sequences were annotated to Eukaryota, and Fungi accounted for 69.55%, Viridiplantae 29.52%, Metazoa 0.82% and unclassified Eukaryota’s sequences 0.11%.

Studies have shown that zokor is a pure herbivorous animal, with a well-developed cecum for decomposing cellulose in plants [50]. Meanwhile, we classified the fungi groups obtained, and found that they were mainly pathogenic fungi of plants, endophytic fungi, fungi in soil, and fungi intrinsic to the stomach of zokor, but not large edible fungi. Metazoan analysis found that the main species were ticks and mites on plants, as well as *Canis*. The above species may be absorbed by the zokor when feeding on the roots and stems of plants, and whether there is active feeding needs further study. In the analysis, we deleted all the above groups and did not do the analysis, which mainly focused on Viridiplantae.

### OTU vs ASV taxonomic analysis

We used the OTU clustering method and the ASV method to make taxonomic annotation on the amplified feeding sequences, and compared the types annotated by the two methods at the Order, Family, Genus and Species levels. At the level of Order, Family and Genus, there is a high degree of coincidence between the annotated information obtained by the two methods, but at the level of Species, the number of overlapped species of the two methods decreases, and the annotated information of the two methods at the level of Species is greatly different. We found that ASV method can improve the diversity of food species than OTU clustering method, that is, more species can be annotated. Therefore, ASV method was selected for subsequent dietary analysis. (**Table 2**.)

**Table 2.** Comparison of annotating ability between OTU clustering method and ASV method.

|  | OTU particular | shared | ASV particular |
| --- | --- | --- | --- |
| Order | 2 | 23 | 2 |
| Family | 5 | 31 | 5 |
| Genus | 29 | 65 | 27 |
| Species | 77 | 73 | 93 |

### Pan/Core species analysis

From the Pan species curve, we can see that the curves of species at Order and Family level have been flat, but at Genus and Species level, the curves are still in an increasing stage, indicating that our sample size cannot fully reflect the food composition of the study area, and further expansion of the sample size is needed in the follow-up studies.(**Fig.1a**) According to the Core species curve, with the increase of sample size, at 4-5 samples, there are Core species at Order and Family level, while the number of Core species at Genus and Species level tends to zero. (**Fig.1b**)

**Fig 1.**
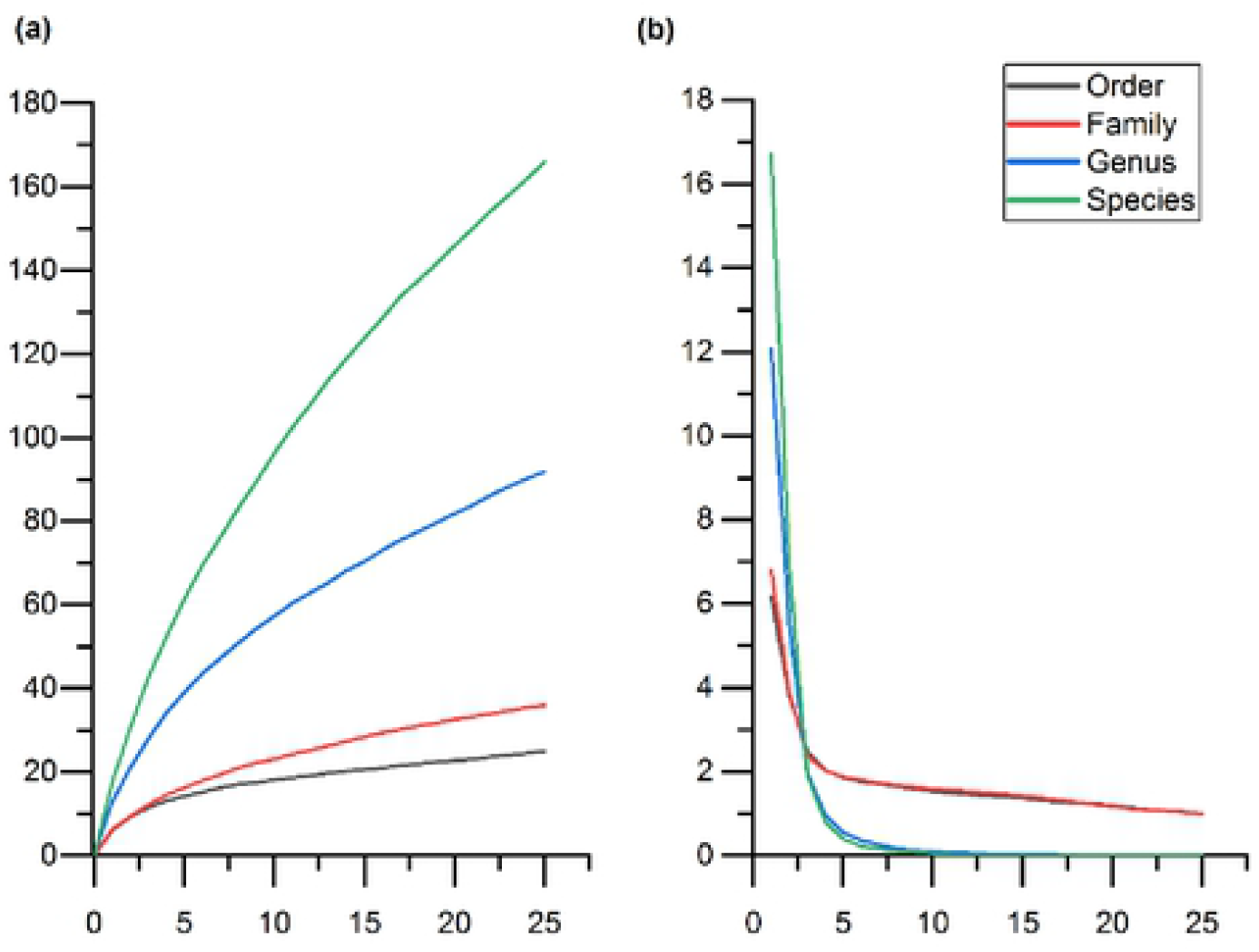
Pan/Core species analysis diagram. **a** and **b** respectively show the Pan and Core species analysis at the Family, Genus and Species levels of plant species after the combination of the two primers. The horizontal axis represents the number of samples. In **a**, the vertical axis represents the number of Pan species and in **b**, the vertical axis represents the number of Core species.

### Venn diagram of dietary of zokor

In the 7 study areas, there were 4 shared food families (Asteraceae, Poaceae, Rosaceae and Convolvulaceae) (**Fig.2a**,**3a**), and 4 shared genera *(Artemisia,Echinops, Cirsium and Convolvulus*) (**Fig.2b**,**3b**). The main plant genera were*Artemisia, Echinops* and *Cirsium*(**3b**). In addition, there were significant differences in food species of zokor in different areas. Combining with plant species investigation in the study area, the differences in food species of zokor in different study areas were consistent with the differences in plant species among plots, which indicated that zokor was a typical opportunist who first fed on plants near the burrow [21].

**Fig 2.**
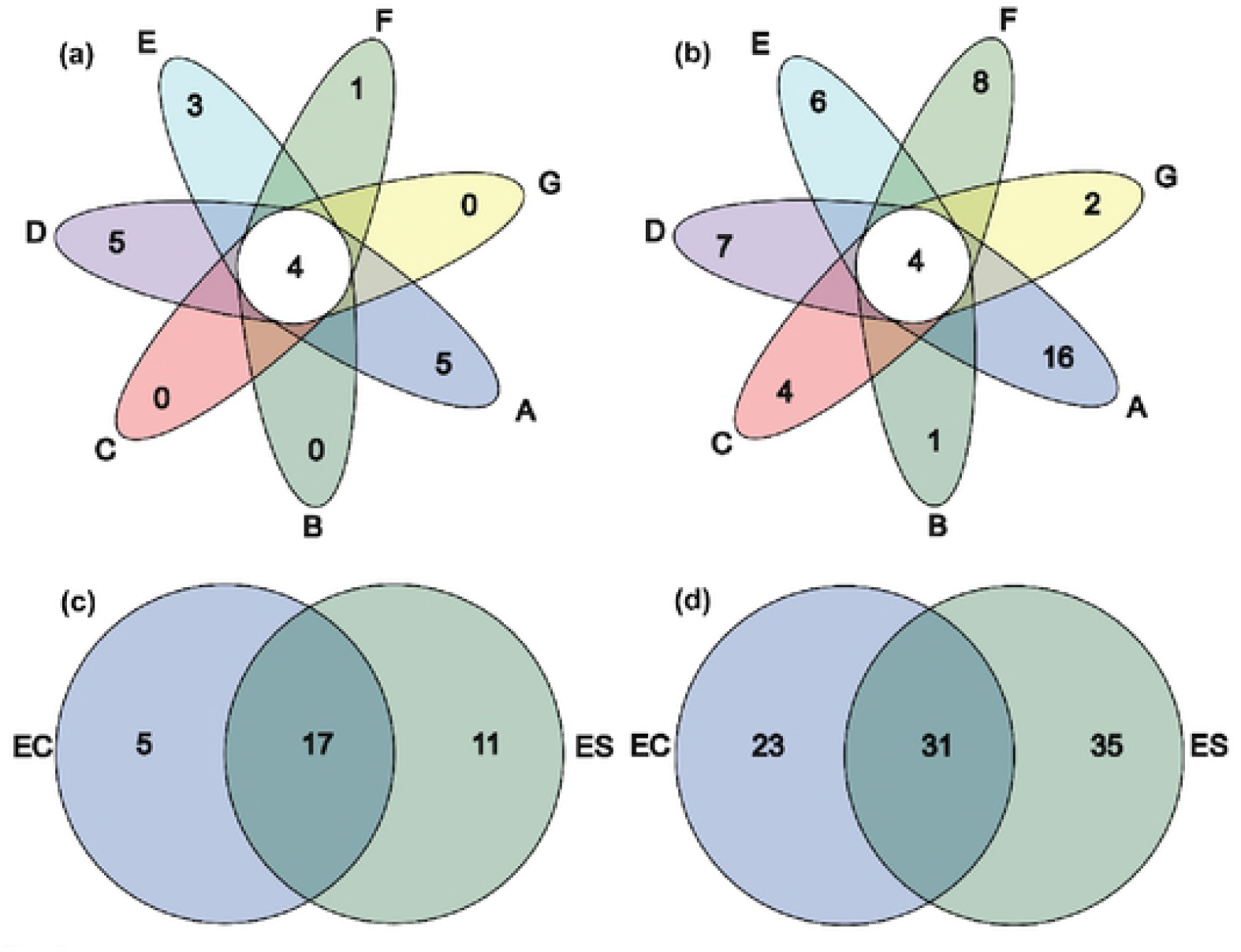
Venn diagram of dietary of zokor. **a** and **b** were the food composition at the Family level and Genus level, respectively,**c** and **d** represent the food composition at the Family and Genus levels, respectively.

**Fig 3.**
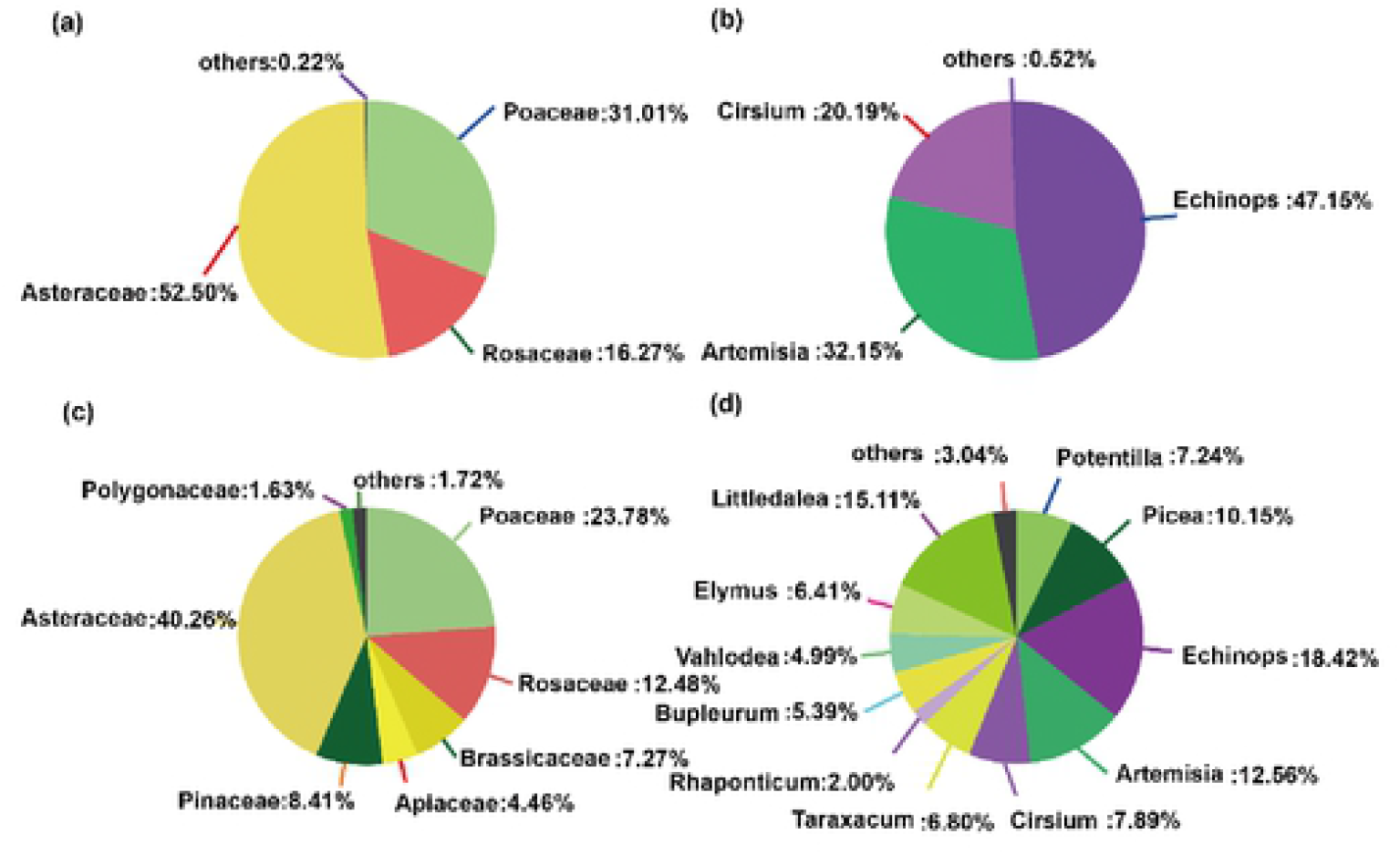
Venn plot shared. **a** and **b** represent the families and genera shared at all areas, respectively,**c** and **d** represent families and genera of plants that these two species eat together.

Among the food composition of the two species of zokor, at the Family level, 17 plant families were feeding for both species of zokor (**Fig.3c**), and at the Genus level, 31 plant genera were feeding for both species ( (**Fig.3d**). The Venn diagram indicated that the dietary overlap of the two species was high, especially at the Family level.

### Diet of zokor after the combination of two primers

After combining the results annotated by the two primers, the total food composition of 25 zokors was obtained. The food composition of zokors was evaluated by %RA and %FOO, respectively. We excluded species with %RA less than 0.01% from the analysis, on the one hand, because their ASV numbers were too low and their %FOO was too low to introduce bias if they were included in the analysis [51], on the other, the importance of rare food categories may be overestimated if the analysis is conducted only by ordering the occurrence.

By combining %RA and %FOO methods, we evaluated the diet of the zokor at the Family and Genus levels. We found that (1) at the Family level, %RA of Asteraceae was 38.16%, while %FOO was 100%, indicating that every zokor fed on Asteraceae. %RA of Poaceae was 22.52%, %FOO was 96%, and only one zokor did not feed on grassy weed. This indicates that the Asteraceae and Poaceae play an extremely important role in the zokor’s diet, next for Rosaceae, Pinaceae, Brassicaceae and so on. The tendency of %RA and %FOO at the Family level showed relatively good consistency. (**Fig.4**);(2) at the genus level, the %RA of *Echinops* is 14.45%, which is the highest among all plant genera, but the %FOO is only 56%. The %RA and %FOO of *Littledalea* are 11.86% and 40% respectively. %RA of *Artemisia* was 9.85%,surprisingly, %FOO was the highest among all plant genera, 84%. In addition, some plant genera with relatively important values of %RA and %FOO include *Cirsium, Elymus, Potentilla, Bupleurum*, etc.(**Fig.5**). The detailed %RA and %FOO tables are shown in the attached tables.(**Appendix 1**)

**Fig 4.**
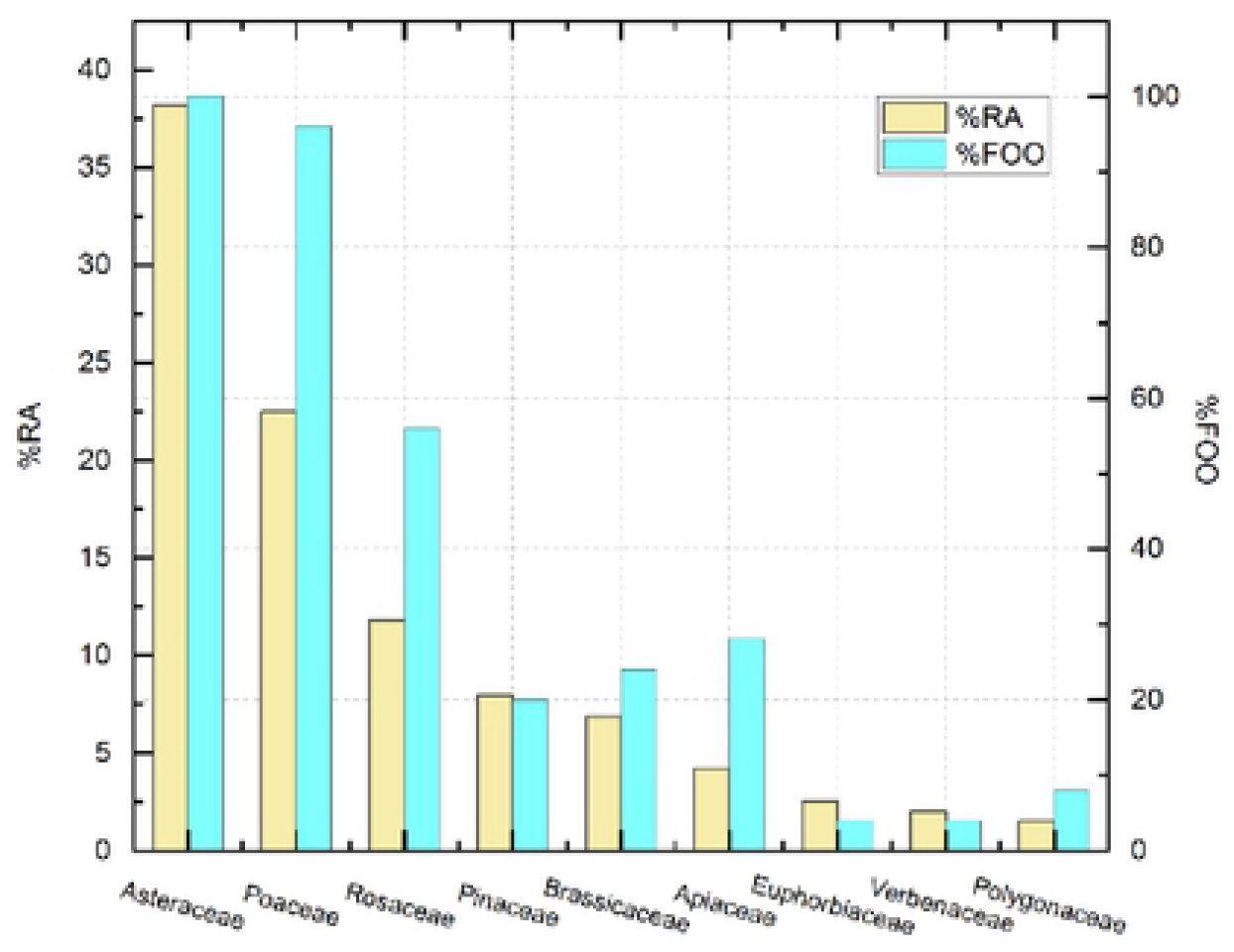
% RA and % FOO represented plant families.

**Fig 5.**
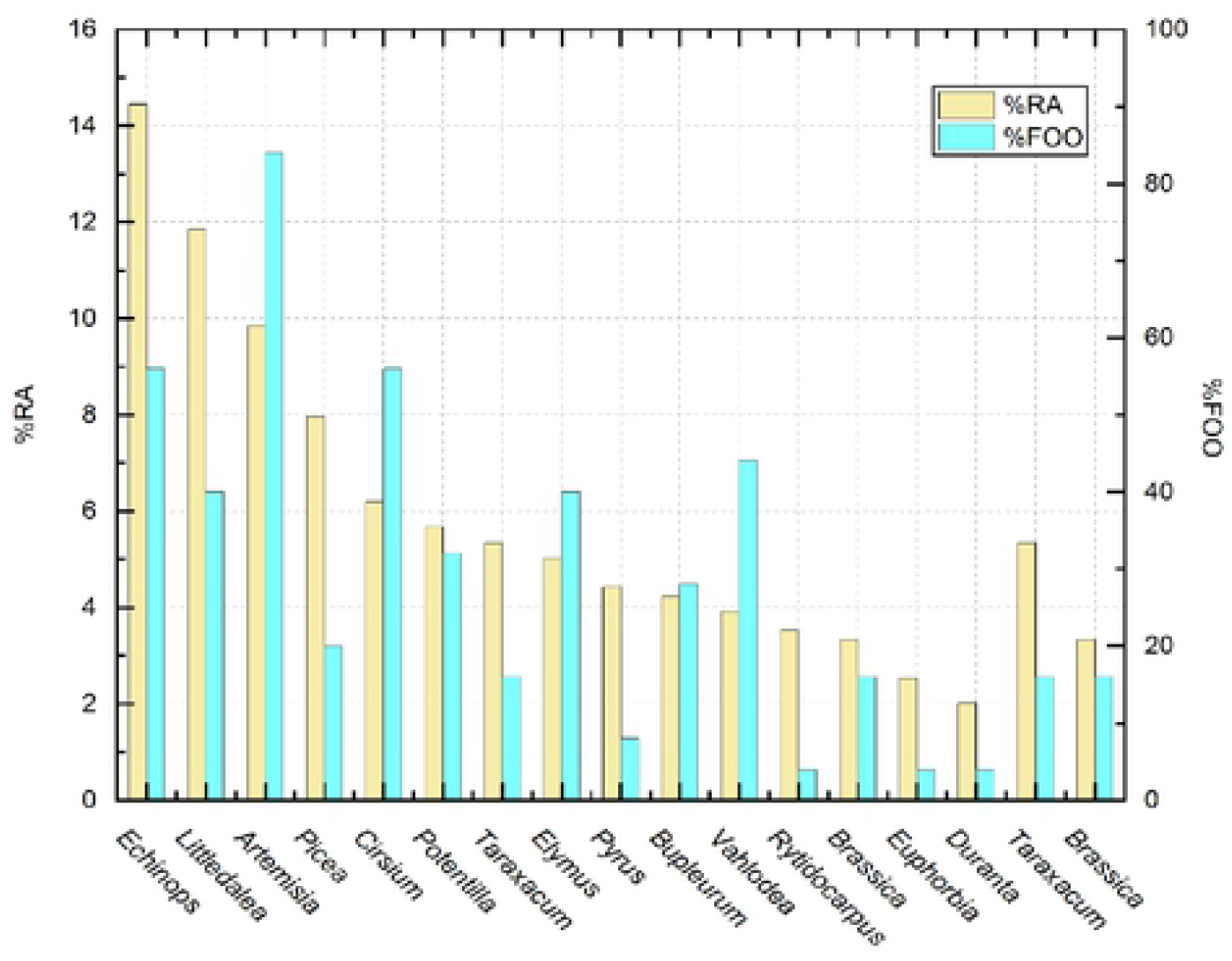
% RA and % FOO represented plant genera.

**Fig.4** and **Fig.5** respectively showed that at the Family level, the zokor mainly fed on plants of Poaceae, Asteraceae and Rosaceae, at the Genus level, the zokor mainly fed on plants of the *Echinops, Artemisia, Picea* and *Cirsium*.**Fig.6** and **Fig.7** respectively showed the diet of zokor at the Family and Genus level by %RA method.

**Fig 6.**
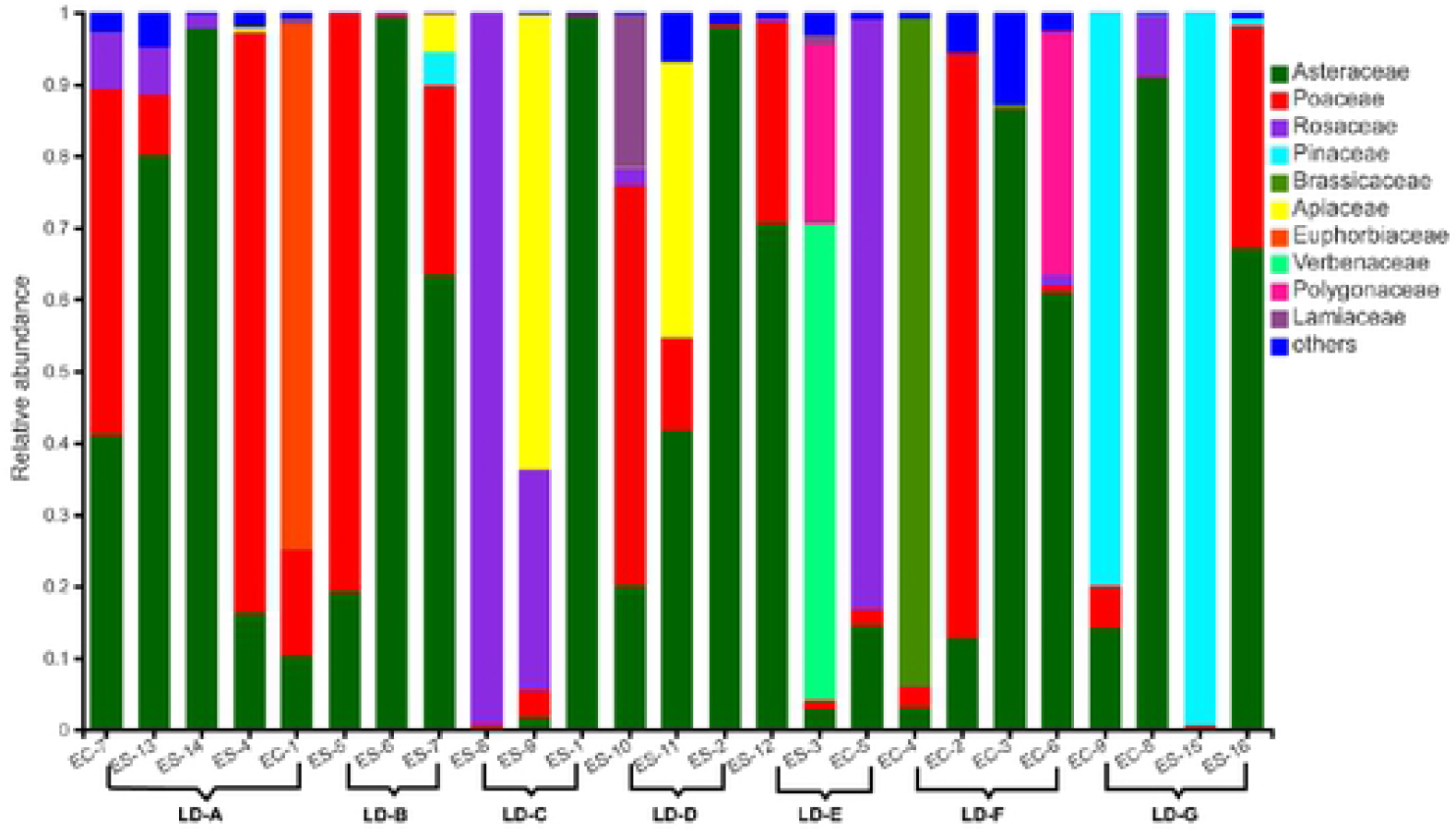
Diet of zokor at Family level. other<0.1,n=25

**Fig 7.**
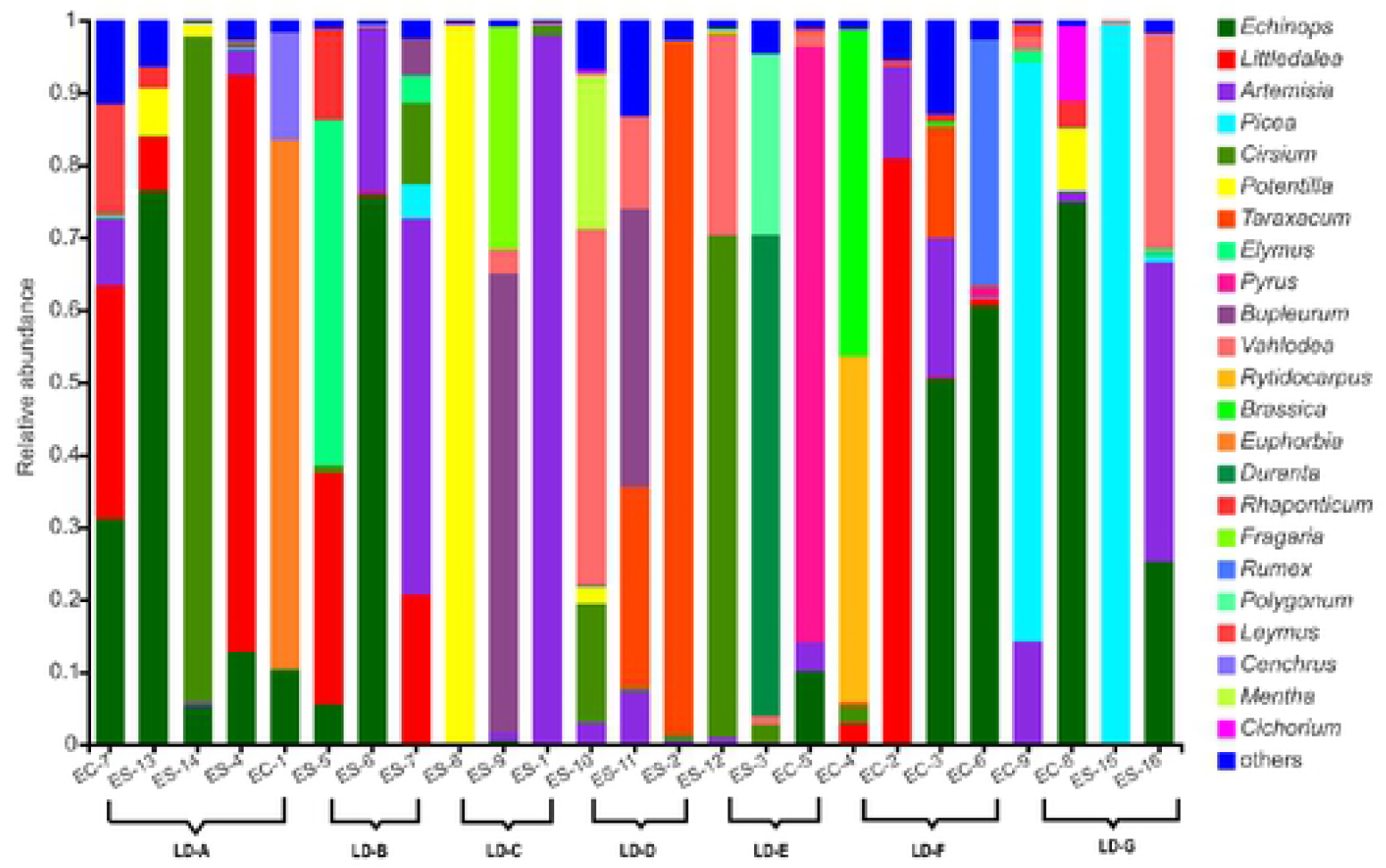
Diet of zokor at Genus level. other<0.1,n=25

### Circos diagram of food composition of zokor

We produced Circos diagram to show the detailed correspondence between the diet of zokor in different groups. **Fig.8** is the correspondence analysis of diet of two species of zokor at the Genus level of food, and **Fig.9** is the correspondence analysis of zokor’s diet at the Genus level in 7 plots.

**Fig 8.**
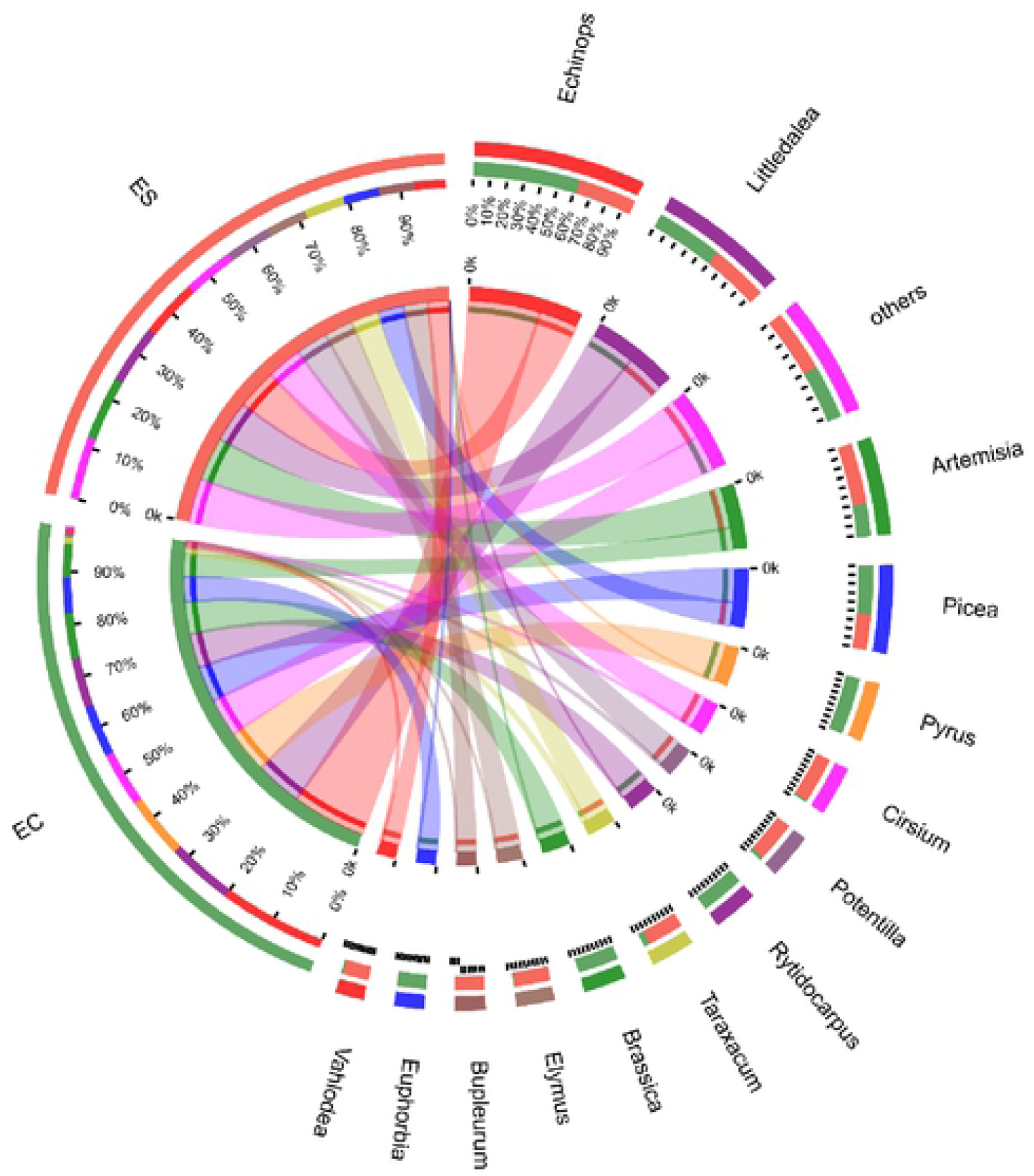
Circos diagram of diet of two species of zokor in the interspecific. Genera with relative abundance less than 5% are classified as others.

**Fig 9.**
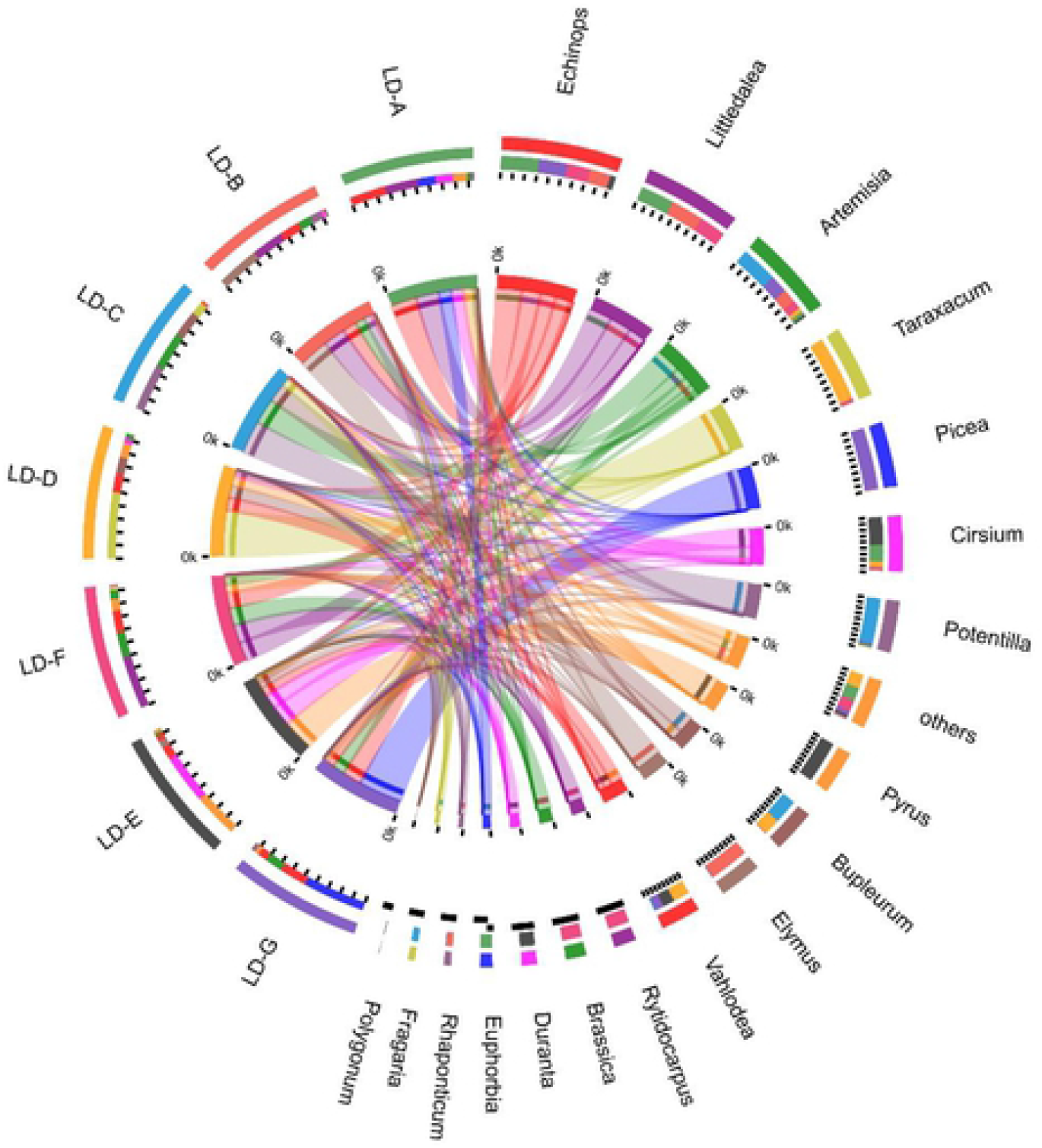
Circos diagram of diet at Genus level of zokor between different study area. Genera with relative abundance less than 5% are classified as others.

### Diversity analysis of zokor’s diet

#### Alpha diversity analysis

As Sobs index, Chao 1 index and Ace index have almost no difference in the calculated diversity value, so only the Sobs index was selected for analysis. Coverage index was always 1 because ASV which only appeared once was removed in the sequence annotation, so the analysis of the Coverage index was not carried out.

At the Family level, the zokor’s diet Sobs index ranged from 3.67-7.25 (mean 5.75), and Shannon ranged from 0.32 to 0.82 (mean 0.54), indicating low diversity. At the Genus level, the zokor’s diet Sobs index ranged from 9.33 to 14.40 with an average value of 11.35, and the Shannon index ranged from 0.36 to 1.10 with an average value of 0.86, indicating that the diversity of diet was low.

The diversity of zokor’s diet differs greatly among different study areas, which may be related to the plant species distributed in different study areas. Shannoneven index reflects the evenness of eating habits of zokor, which has the same trend as the Shannon index. In the study area with a high Shannon index, the evenness of food species is also high.(**Table 3**.)

**Table 3.** Alpha diversity index table.

|  | Family |  |  | Genus |  |  |
| --- | --- | --- | --- | --- | --- | --- |
|  | Sobs | Shannon | Shannoneven | Sobs | Shannon | Shannoneven |
| LD-A | 6.80±0.98 | 0.67±0.31 | 0.34±0.16 | 14.40±4.45 | 0.92±0.43 | 0.34±0.14 |
| LD-B | 3.67±0.94 | 0.50±0.37 | 0.36±0.23 | 11.00±2.16 | 1.10±0.36 | 0.46±0.12 |
| LD-C | 5.00±1.63 | 0.32±0.39 | 0.17±0.19 | 9.33±2.62 | 0.36±0.38 | 0.15±0.14 |
| LD-D | 6.67±1.25 | 0.82±0.49 | 0.41±0.24 | 13.33±2.62 | 1.10±0.60 | 0.43±0.24 |
| LD-E | 6.33±1.25 | 0.73±0.16 | 0.40±0.06 | 10.00±1.41 | 0.80±0.12 | 0.35±0.05 |
| LD-F | 7.25±0.43 | 0.59±0.19 | 0.30±0.10 | 11.75±2.77 | 1.00±0.28 | 0.41±0.09 |
| LD-G | 4.50±1.66 | 0.43±0.27 | 0.31±0.21 | 9.75±3.70 | 0.73±0.45 | 0.30±0.17 |
| EC | 6.56±1.34 | 0.64±0.23 | 0.36±0.14 | 12.00±3.33 | 0.98±0.33 | 0.39±0.10 |
| ES | 5.44±1.80 | 0.55±0.41 | 0.31±0.22 | 11.31±3.79 | 0.80±0.51 | 0.32±0.20 |

In comparison of food species between *E. smithii* and *E. cansus*, the food diversity of *E. cansus* was slightly higher than that of *E. smithii* in terms of Sobs index, Shannon index and Shannoneven, but the sample size was smaller than in *E. smithii*. (9<16). Zokor was the dominant species in this study area. Our results suggest that the *E. cansus* avoids interspecific competition with its relative *E. smithii* by adopting a more diverse diet. Due to its large population, the *E. smithii* consumes more available food resources in this area.

#### Difference test between groups of indices

Kruskal-Wallis rank sum test and FDR multiple test correction were used to test the difference between index groups, and there was no significant difference between Family and Genus and among different species of zokor (*P*>0.05) (data not shown).

#### Correlation analysis of diet alpha diversity indexes and body indexes

We measured the body indexes of zokor, including body weight and body length, and used Pearson double-tailed significance test to find the correlation between diet diversity index (Sobs index, Shannon index, Shannoneven index) and body indexes. The results showed that the body weight and body length of zokor were slightly negatively correlated with the alpha diversity indexes, and there was no significant difference. The correlation coefficient between Sobs index and body weight was −0.382 (*P*=0.059). Our results seem to indicate that diet diversity may decrease with greater body size. Larger zokor are better at obtaining food than smaller ones, so the variety of food available may be relatively simple.

#### Beta diversity analysis

##### Principal component analysis(PCA)

PCA results showed that almost all (22/25) of the gastric content samples were clustered together, with only EC-7, EC-4 and ES-5 deviating from most of the samples. (**See Fig S2**) These results may indicate that (1) the plant species in our study area may have low heterogeneity, and the zokor’s diet in different study areas was mostly clustered together; (2) The interspecific competition between the two zokor species may not be strong, and the food between the two species is similar.

### Analysis of food selectivity

Through investigating the research area of the plant resources, form a list of plants in the study area (**Appendix 2**), compare the zokor feeding selectivity index of sample area plant selection in the sample area coverage of more than 0.1% of the plant, comparative DNA metabarcoding on the plants of the Genus of zokor feeding ratio, selectivity index calculation. Ivlev’s [52] selectivity index was used to calculate the plant selectivity of zokor. E_i_ = (r_i_-p_i_)/(r_i_ + p_i_); r_i_: refers to the proportion of i plants in the feeding composition. p_i_: refers to the proportion of i plant biomass to the total biomass in the plant survey quadrate. E_i_ is between −1 and +1, if E; >0 indicates that the animal has a positive choice for the plant. E_i_ <0 means negative selection, and E_i_ = −1 means no selection.

It can be seen that the zokor feeds positively on the genera *Echinops, Bupleurum, Crisium, Brassica, Calamagrostis, Medicago, Sanguisorba* and other Poaceae, which account for 21.87% of the plant resources in this area, but 53.22% in the food of the zokor. For the genera *Elymus, Leymus, Artemisia, Poa, Potentilla*, and *Convolvulus* they were widely distributed in this area, accounting for 74.15% of the plant resources, but only 16.43% of the zokor’s diet.(**Fig.10**)

**Fig 10.**
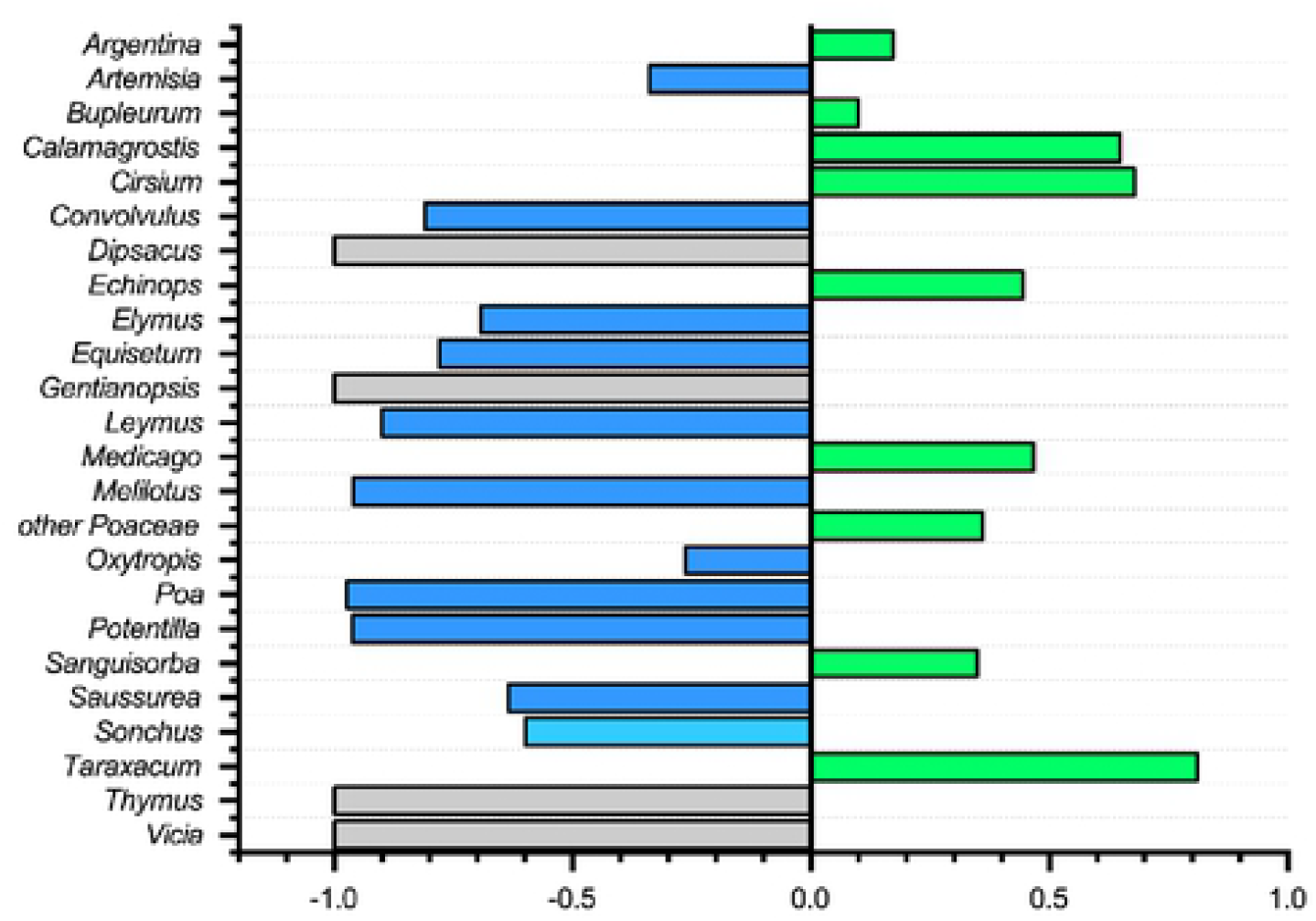
Food selection diagram of zokor. **Green** represents positive selection; **Blue** represents negative selection; **Grey** means do not select.

### Food type analysis

Through the comparison of the annotated food list of zokor by referring to the *Flora of China* and the stored roots dug out from the caves of zokor, the plants fed by zokor can be divided into the following types:

#### Plants are classified in terms of their life forms

Based on the life form of the plant and the abundance of that type of plant in the study area, they can be divided into annual herb (AH), perennial herb (PH), arbor (A), and other types. Other types include shrub (S), biennial herb (BH), floating algae (FA) and fern (F).(**Fig.11**)

**Fig 11.**
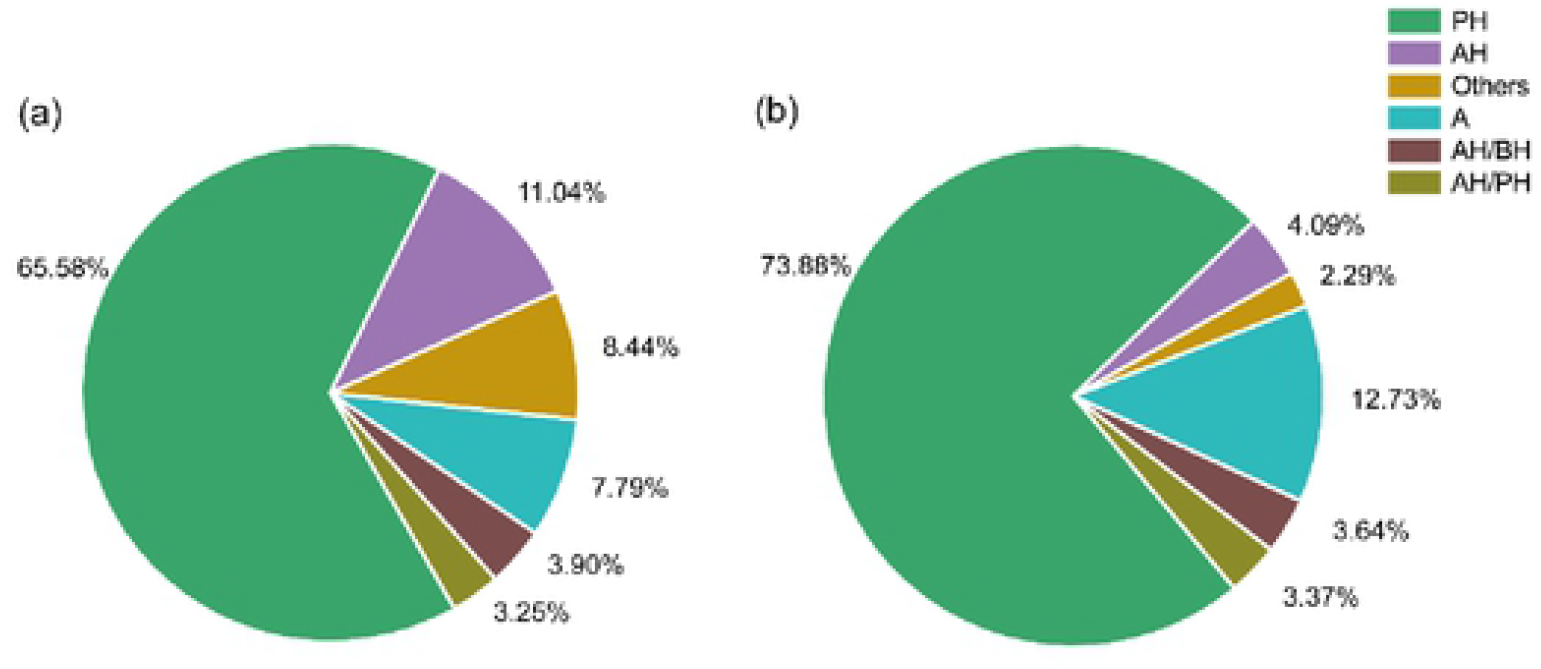
Pie chart of plant feeding parts of the zokor. **a** and **b** were the proportion of the number of plant species and the number of ASV of the feeding type of zokor.

#### Plants are classified in terms of their feeding parts

We briefly discussed and analyzed various types of food according to the plant parts that zokor feeds on. The rat mainly feeds on taproots(54.60%) and rhizomes(26.41%),(**Table 4**.) which are rich in water and energy, which can meet the high energy consumption caused by zokor’s digging habit [50].

**Table 4.** Analysis of the feeding parts of plants.

| foraging types | root type | representative | features | %RA |
| --- | --- | --- | --- | --- |
| tree root | coniferous tree | *Picea asperata* | fibrous and fatty [55] | 7.99% |
|  | other tree | *Pyrus,Diospyros* | rich in fiber | 6.77% |
| herbaceous root | rhizome | *Elymus,Artemisia* | reproductive part [54–56] | 26.41% |
|  | taproot | *Bupleurum,Taraxacum,* | thick,watery and nutritious [57] | 54.60% |
|  | tuber&bulbus | *Stachys geobombycis* | watery and sugary [58] | 0.23% |
|  | fibrous root | *Setaria viridis* | / | 3.93% |
| other | rootless | FA,F | / | 0.06% |

## Discussion

In this study, the zokor’s diet endemic to China was determined by DNA metabarcoding based on HTS, which was the first report on the food composition of *E. smithii*. Our results suggest that DNA metabarcoding is feasible for studying the feeding habits of rodents that live entirely underground and feed mainly on plant roots. In the study area, *E. smithii* is sympatric with *E. cansus*, and both belong to *Eosplax*. Studies have shown that the diet of seven species of *Ctenomys* in Brazil has a high overlap [4].Our results also confirmed that there was no significant difference in food composition between *E. smithii* and *E. cansus*, with schoener niche overlap index of 0.69. In this area, two species of zokor mainly fed on the roots of PH, which mainly belonged to genus *Echinops, Artemisia, Cirsium, Bupleurum* and *Elymus*, and also fed on the roots of *Picea* and *Pyrus*. Since the samples we selected were the gastric contents of zokor and ITS primers were used to amplify the DNA of the gastric contents, many fungi were also amplified because the digestion degree of food in the stomach of mammals was the lowest, but the main groups were common plant pathological fungi, endophytic fungi, intestinal fungi, etc. (data not shown). It’s either eaten by the rat as it feeds on the roots, or it’s already present in the stomach, which may not be the result of the zokor’s choice of food. This was also the case in the metabarcode feeding studies of lemmings’ stomach contents [28].

Interestingly, ITS also amplified *Canis*. Since there were no stray dogs distributed in the study area, it was preliminarily speculated that it might be *Canis lupus*, which may be that zokor ingested wolf feces in the process of digging and eating. The common sense now is that birds of prey (Falconiformes,Strigiformes) and small predators (Canidae, Mustelidae,Felidae) are natural enemies of the zokor, and it’s easy to understand that they are, and the zokor is also a rat. But this view is one-sided, zokor rarely appears on the ground. Its temporary herbivorous burrow is 10-30cm from the ground, and its permanent cave is 50-210cm from the ground [13]. Whereas raptors, who hover high in the air and use their keen vision to spot their prey, apparently couldn’t spot the zokor, there is no study or report on prey of zokor by raptors. Small, slender carnivores, such as the *Mustela sibirica*, that can burrow into the zokor’s hole to prey on the zokor, are more plausible, but research on this is lacking. Further research is needed to determine whether the wolf we have found indicates that it is a natural enemy of the zokor.

And we found something interesting in the species of plants that we annotated, Hydrodictyaceae(EC-2,ES-11,EC-6), Polypodiaceae (ES-2), these plant species are unique in that they will only live where there is watery or humid. Streams in the study area confirmed the possibility of the Hydrodictyaceae. Both EC-2 and EC-6 belong to LD-F plot, while ES-11 and ES-2 belong to LD-D plot, and their distance from streams is 180m and 600m, respectively. This suggests that the zokor is adapted to moist conditions, and that it travels considerable distances in search of food.

By analyzing the zokor’s diet and comparing the plant types in the study area, we found that zokor is a typical opportunist, that is, it feeds on most of the plants in the study area, which is consistent with many studies on the feeding habits of herbivores [4,21,24,26]. Since most of the *Picea, Pinus mongolicus* and *Pinus tabulaeformis* in the study area have carried out wire nets in their roots, which can effectively prevent the biting of zokor, they then turned to feed on the rhizome of PH such as Poaceae and Asteraceae widely distributed in the area. The zokor’s food storage showed that almost entirely the asexual part of the plant.

It was autumn when we caught the rat, which is the season for storing food. We also found two storehouses, one of which had five small pantries, and the other only three small pantries, which were neatly stuffed with plant roots. It can be mainly divided into gramineous weeds and forbs. Gramineous weeds account for 28.18%-34.23% of raw weight, and forbs account for 64.84%-69.74%. Different from the *E. baileyi* [21], the zokor warehouse we found contains only the hypogeal parts of plants, but not the aerial parts. It has been found that the zokor spends part of its time on the ground, which may allow it to feed on the aboveground parts of plants [59]. In autumn, the density of *Ctenomys* was positively correlated with the availability of plants with reserve organs [60]. Similarly, the zokor becomes significantly more active in autumn, actively searching for plant reserve organs (rhizomes, tubers, taproots, etc.). In food selection, taproot (54.60%) and rhizome (26.41%) were the main food types, and these types of roots provided water, carbohydrate, fat and protein for zokor. For example, dandelion roots contain 22.59% more carbohydrates than aerial parts [57]. The root tuber of *Stachys geobombycis* contains 23g of sugar, 4.1g of protein, 0.3g of fat and nearly 72% of water per 100g [58].

Studies have shown that grassland management such as grazing and mowing to reduce litter accumulation can alleviate the negative impact of nitrogen deposition on plant diversity by reducing the asexual reproduction of dominant species [56]. Our study area is dominated by PH, and the zokor’s feeding on the asexual reproductive organs of these herbaceous plants may reduce the asexual reproduction of dominant species and indirectly slow down the decrease of plant diversity. In our study area, and in a larger area of woodland, the zokor is considered a pest mainly because it gnaws on the roots of afforestation plants. The contribution of woodlands to soil and water conservation is well known, however, the role of the zokor in the ecosystem is little-known. Only when the density of zokor was too high, it was forced by intraspecific and interspecific competition to cause harm to forest. Most of the tree species in the study area were covered with wire nets, and our results and field survey showed that zokor did little harm to these trees. According to the literature, zokor can optimize the composition of meadow community with appropriate density [22]. At the same time, as the basic species in the food web, zokor plays a positive role in maintaining the stability of the ecosystem and promoting energy flow. We should not destroy zokor blindly and unilaterally. Our results showed that the zokor did not cause great harm to the tree species in the study area. It is not that the zokor’s habits have changed,outside the study area, there are still many pines that were eaten to death. In the subsequent afforestation, we should set up the idea of protecting the target species and tolerate the existence of zokor to a certain extent, which is a win-win situation for the development of forestry economy and ecological restoration.

At present, most studies on diet based on HTS still choose OTU clustering method for sequence annotation [4, 18, 20, 24, 26–28]. We found that the ASV method was similar to the OTU clustering method in the identification ability at the Order, Family and Genus levels, but at the Species level, ASV method had 16 more species than OTU. As with fungal and microbial studies [35, 36], the ASV method also improves the diversity of herbivore analysis.

Our study was compared with that of other researchers on the diet of zokor (**Table 5**). In this study, 65 species of plants belonging to 56 genera and 24 families were investigated in the study area, and 154 species of plants belonging to 80 genera and 32 families were detected by DNA metabarcoding technology.(**Appendix 3**) It can be directly found that DNA metabarcoding can greatly improve the identification of food species and improve the understanding of zokor’s diet. There may be two reasons why zokor fed more species than plants in the study area: (1) As a typical burrowing animal, zokor mainly feeds on the roots of plants, possibly taking in some soil in the process. In the study on the feeding habits of *Ctenomys*, it was found that 23 plant families and 58 MOTUS could be recovered by extracting DNA from soil alone, and the diversity of plant families and MOTUS in soil samples was higher than that in the feces of *Ctenomys* [4]. Therefore, there may be false positives in our experimental results, that is, the actual variety of food resources consumed may be smaller than that recovered by DNA metabarcoding.(2) The time when we investigated the plant species in the study area was autumn, when vegetation and trees were dying, and some species were not investigated.

**Table 5.** Comparison of diet of zokor.

| researchers | species | method | specimens number | resources number | feeding number |
| --- | --- | --- | --- | --- | --- |
| Wang QY et al. [22] | *E. baileyi* | microhistology | 92 | 67 | 28 |
| Su JH et al. [23] | *E. baileyi* | microhistology | 40 | 18 | 11 |
| Xie JX et al. [56] | *E. baileyi* | dig the zokor’s pantry | 57 | 66 | 59 |
| Wang J [16] | *E. cansus* | captive feeding | 16 | 74 | 71 |
| Bazhenov YA [62] | *M. aspalax* | dig the zokor’s pantry | 3 | / | 11 |
| this study | *E. cansus* & *E. smithii* | DNA metabarcoding | 25 | 65 | 154 |

Indeed, in the Pan species graph, our specimen has not flattened the observed species curve, which requires more samples to be added for food analysis of zokor. In our study area of 300 ha with similar habitat types, we need to collect samples over a large geographical range and a long time period to avoid bias [63]. Fortunately, our samples were not collected at the same time or within a few days. Our sampling time span was nearly two months. In general, larger sample sizes and technical repetitions of a single sample (multiple extraction, multiple sequencing) are certainly beneficial [64, 65]. However, due to the increasing workload and cost, researchers cannot continuously increase the sample size and increase the repetition. As a result, they need to find the best balance between sample size and biological and technical replication, taking into account the workload and cost [63]. The results of our study are credible at the level of Genus classification and can explain the dietary characteristics of zokor under natural conditions. Many herbivorous animal feeding studies are also conducted at the level of Family [30]and Genus [26, 66]. In the study of feeding habits of DNA sequencing, there may be deviations/errors in every link, such as sample preservation, sample contamination, DNA extraction, PCR amplification, primer selection, library preparation, selection of sequencing platform, error removal, sequence taxonomic assignment, etc., which will affect the final result. Even so, the classification accuracy of diet studies based on HTS is much higher than that of traditional microscopical methods, and the workload is much less than that of microscopical methods.

## Supporting information

**Fig S1.**
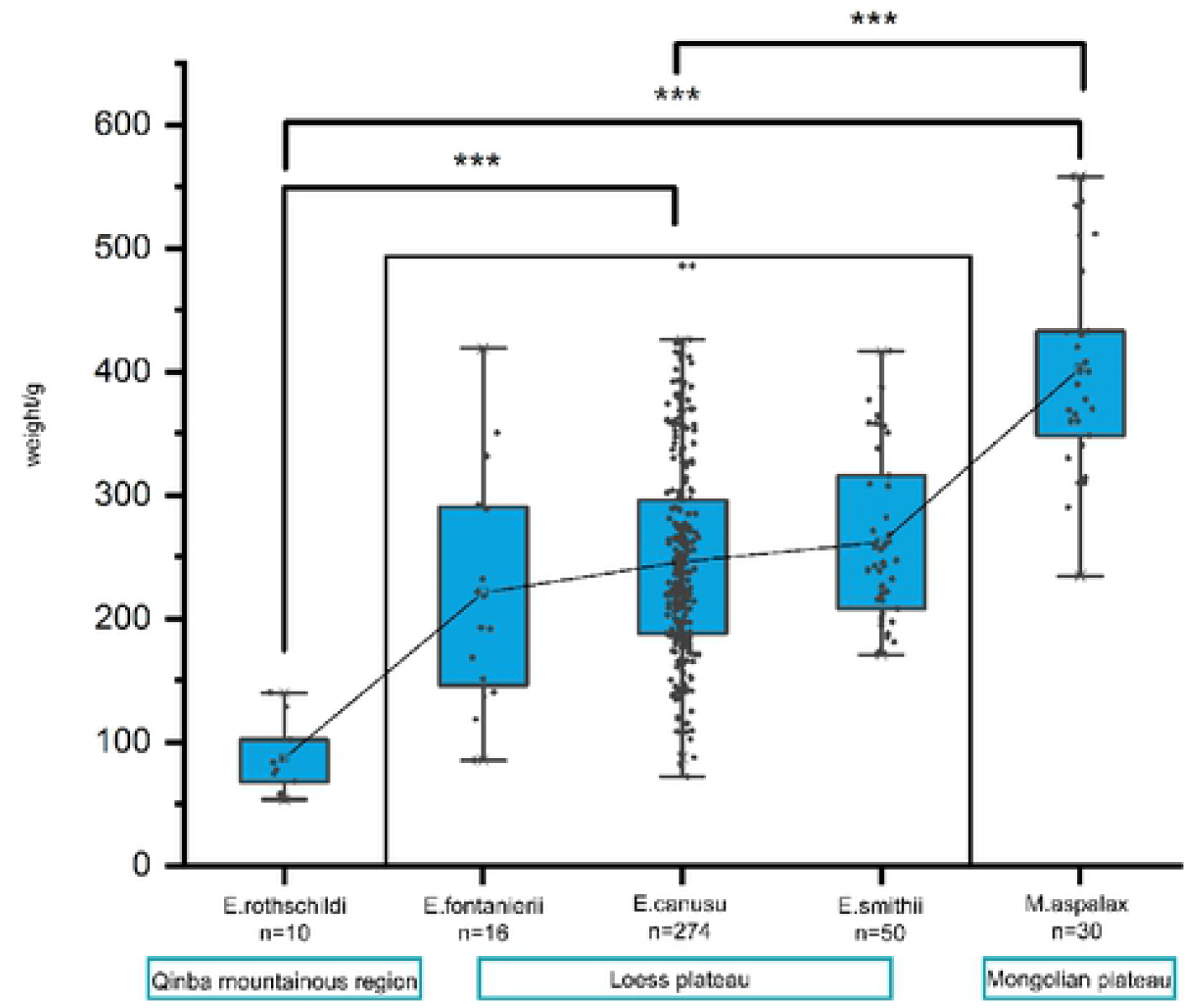
Body weight of zokor in different geographical regions. (TIFF)

**Appendix 1 Detailed %RA and %FOO tables.**

(EXCEL)

**Appendix 2 Flora of study area.**

(EXCEL)

**Appendix 3 Detailed DNA metabarcoding diet tables.**

(EXCEL)

**Fig S2.**
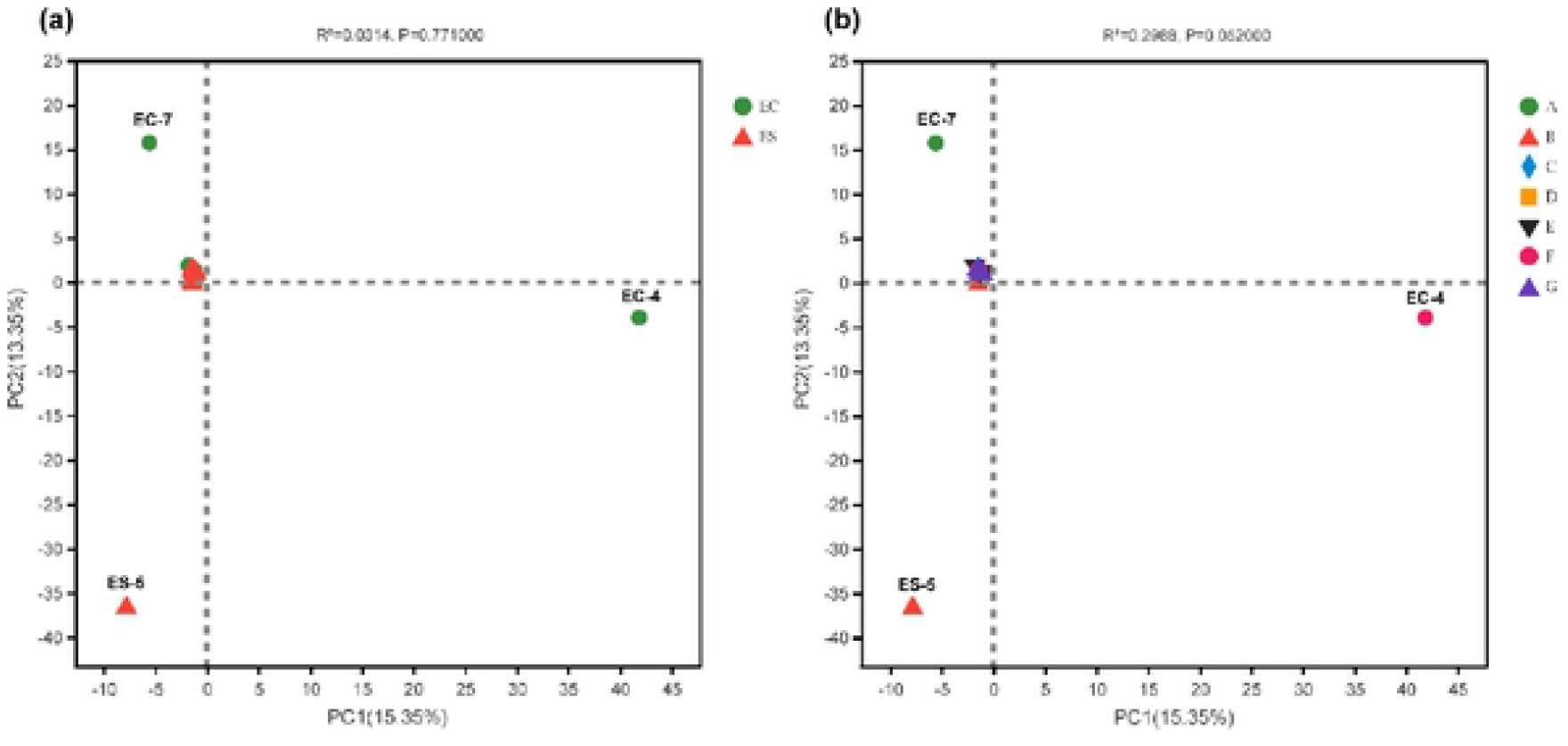
PCA plot. **a** represents the PCA analysis of two species of zokor, and **b** represents the PCA analysis of 7 sample plots (TIFF)

## Acknowledgments

We sincerely thank the Forestry Pest Control Station of Longde County, Ningxia Hui Autonomous Region for its assistance in the collection of zokor samples.

All authors contributed to the study design. Sample collection and data analysis were performed by Xuxin Zhang,Yao Zou,Xiaoning Nan and Chongxuan Han. The first draft of the manuscript was written by Xuxin Zhang. Experiment was performed by Xuxin Zhang, Yao Zou.The revision suggestion of this paper was proposed by Chongxuan Han and Xiaoning Nan.All authors have read and approved the final manuscript.

## Notes

### Competing Interest Statement

The authors have declared no competing interest.

